# Nef and Vpu protect infected cells from ADCP mediated by plasma from people with HIV-1

**DOI:** 10.64898/2026.05.14.725178

**Authors:** Étienne Bélanger, Alexandra Tauzin, Fitsumbrhan Tajebe, Monika Chandravanshi, Derek Yang, Hung-Ching Chen, Ta-Jung Chiu, Catherine Bourassa, Halima Medjahed, William D. Tolbert, Madeleine Durand, Jonathan Richard, Donna M. Huryn, Simona Stäger, Marzena Pazgier, Andrés Finzi

## Abstract

The HIV-1 envelope glycoprotein (Env) represents the only viral antigen at the surface of infected cells, making it an ideal target for antibody-based therapies. Most antibodies elicited in people with HIV (PWH) do not recognize Env in its native “closed” conformation but readily bind to Env when it samples the CD4-bound “open” conformation. Downregulation of CD4 at the surface of infected cells by the viral accessory proteins Nef and Vpu prevents the premature opening of Env and has been shown to protect infected cells from antibody-dependent cellular cytotoxicity (ADCC) mediated by PWH plasma. Here, we report that deletion of Nef and Vpu from primary infectious molecular clones renders infected cells vulnerable to antibody-dependent cellular phagocytosis (ADCP) mediated by PWH plasma. This is in part linked to the premature engagement of Env with CD4. In agreement with an “open” Env being vulnerable to ADCP, small CD4 mimetic compounds (CD4mc) sensitize *in vitro*-infected cells and *ex vivo*-expanded CD4 T cells to ADCP mediated by autologous monocytes in presence of PWH plasma. This effect was further improved by increasing cell surface Env through IFN-induced BST-2 upregulation.

**IMPORTANCE:** Developing new therapies to eliminate HIV-1-infected cells is essential to decrease the size of the HIV-1 reservoir. Fc-effector functions such as antibody dependent cellular cytotoxicity (ADCC) have shown potential in eliminating *in vitro-*infected cells and *ex vivo-*expanded infected cells from people with HIV, decreasing the size of the reservoir and delaying viral rebound in humanized mice. Here, we report that antibody dependent cellular phagocytosis (ADCP) can also be harnessed to eliminate HIV-1-infected cells. We show that infected cells harboring “open” Env conformations are susceptible to ADCP-mediated killing in the presence of plasma from people with HIV. A better understanding of the contribution of different Fc-effector functions in the elimination of infected cells could help guide the development of new therapeutic approaches toward an HIV-1 cure.

## INTRODUCTION

The HIV-1 envelope glycoprotein (Env) is the only viral antigen expressed at the surface of infected cells, making it a prime target for antibody-based therapeutics. Env is synthesized as a gp160 precursor, which is trimerized, glycosylated and proteolytically cleaved during its trafficking to the cell surface, where it is expressed as a trimer of gp120 and gp41 heterodimers [1–3]. Env preferentially adopts a “closed” pre-triggered conformation prior to interaction with its receptor, CD4 [4]. Interaction with CD4 triggers a series of conformational changes in Env eventually leading to the adoption of a fully “open” conformation which enables viral fusion [4]. The vast majority of antibodies elicited in PWH are deemed non-neutralizing (nnAbs), due to their inability to recognize Env in its “closed” conformation and neutralize viral particles [5]. However, the premature interaction of Env with CD4 leads to the exposure of vulnerable Env epitopes which are otherwise occluded in the pre-triggered Env conformation [6]. These epitopes are readily recognized by nnAbs in plasma from people with HIV (PWH) resulting in the elimination of infected cells by antibody-dependent cellular cytotoxicity (ADCC) [7]. To counteract this, HIV-1 has evolved multiple mechanisms to downregulate CD4, notably through the action of its accessory proteins Nef and Vpu [8, 9], but also through mechanisms mediated by Env itself [10–12]. Small CD4 mimetic compounds (CD4mc) bind to the conserved gp120 Phe43 cavity and induce similar Env conformational changes than CD4 [13, 14] resulting in the exposure of vulnerable CD4i epitopes and sensitization of HIV-1-infected cells to ADCC mediated by CD4i Abs and PWH plasma [15–20].

Antibody-dependent cellular phagocytosis (ADCP) is an additional Fc-effector function which occurs when antibody-bound cells are recognized and phagocytosed by effector cells such as monocytes or macrophages. Phagocytes can mediate ADCP via different Fc receptors: FcγRIIa (CD32) and FcγRI (CD64) in the case of IgG-mediated responses and FcαRI (CD89) in the case of IgA-mediated responses [21–23]. ADCP exerts a protective role during infection with pathogens such as Influenza and SARS-CoV-2 [24], however its role in HIV-1 is less clear. There have been reports that various Env-targeting broadly neutralizing antibodies (bNAbs) such as 10-1074, 3BNC117 and 2F5 could mediate ADCP of HIV-1-infected cells *in vitro* [25, 26]. One study, performed on a cohort of more than 500 viremic PWH, reported that participants who were homozygous for the low affinity FcγRIIa allele had a faster disease progression compared to those that were heterozygous or homozygous for the high affinity allele [27], suggesting that ADCP could be involved in the control of HIV infection. Additionally, retrospective studies performed on participants from HIV-1 vaccine trials have linked the ability of antibodies to mediate ADCP with protection from HIV-1 acquisition [28, 29]. Similarly, vaccinal studies in non-human primates recently reported an association between ADCP responses and protection from SHIV acquisition [30]. However, a caveat to many of these studies is that ADCP activity was inferred through the phagocytosis of beads coated with recombinant Env proteins, which does not recapitulate physiologically relevant Env conformations that would be encountered by the immune system during infection [25].

Here, we developed a biologically relevant FACS-based ADCP assay, where Env is expressed at the surface of HIV-1-infected primary CD4^+^ T cells, using autologous monocytes as effector cells. We show that CD4 downregulation by Nef and Vpu protects infected cells from ADCP mediated by PWH plasma by preventing Env from adopting downstream “open” conformations. We further demonstrate that inducing Env “opening” with CD4mc sensitizes these cells to ADCP mediated by PWH plasma, and this effect can be further enhanced by increasing cell surface Env levels through IFN-induced BST-2 upregulation.

## MATERIAL AND METHODS

The materials and methods used here have been reported in previous studies [11, 17–19, 31] and are summarized below.

### Ethics statement

Written informed consent was obtained from all study participants and research adhered to the ethical guidelines of CRCHUM and was reviewed and approved by the CRCHUM Institutional Review Board (Ethics Committee approval nos. 22.173, 11.063, MP-02-2024-11734 and 11.062). Research adhered to the standards indicated by the Declaration of Helsinki. All participants were adult and provided informed written consent prior to enrollment in accordance with Institutional Review Board approval.

### Cell culture and isolation of primary cells

All cells were maintained at 37°C under 5% CO_2_. HEK293T human embryonic kidney cells (ATCC) were grown in Dulbecco’s Modified Eagle Medium (DMEM, Wisent) supplemented with 5% fetal bovine serum (FBS, VWR) and 100U/ml penicillin-streptomycin (Wisent). Human peripheral blood mononuclear cells (PBMCs) from people with and without HIV-1 (Information on PBMC donors can be found in **Tables S1** and **S2**) were obtained by leukapheresis and isolated by Ficoll density gradient. Cells were cryopreserved in liquid nitrogen until further use. Primary CD14^+^ CD16^-^monocytes, used as effector cells for the ADCP assay (**Figure S1**), were isolated from cryopreserved PBMCs by negative selection using immunomagnetic beads as indicated by the manufacturer’s protocol (StemCell Technologies, Cat#19359) and were maintained in RPMI 1640 (Thermo Fisher Scientific) supplemented with 20% FBS and 100U/ml penicillin-streptomycin. Primary CD4^+^ T cells were isolated by negative selection using immunomagnetic beads following the manufacturer’s protocol (StemCell Technologies, Cat#17952). Upon isolation, CD4^+^ T cells were activated with phytohemagglutinin-L (PHA-L, 10µg/ml) for 48h hours and were subsequently maintained in RPMI 1640 supplemented with 20% FBS, 100 U/ penicillin/streptomycin and recombinant IL-2 (rIL-2, 100 U/ml).

### Antibody production and purification

FreeStyle 293F cells (Thermo Fisher Scientific) were maintained in FreeStyle 293F medium (Thermo Fisher Scientific) at a concentration of 1 x 10^6^ cells/ml at 37°C under 8% CO_2_ with 150rpm agitation. Cells were transfected with plasmids expressing the heavy and light chain of A32, and 17b antibodies (kindly provided by James Robinson), as well as 246D and 2G12 antibodies (NIH AIDS Reagent Program) using the ExpiFectamine transfection reagent following the manufacturer’s protocol (Thermo Fisher Scientific). Seven days later, cells were pelleted and discarded, and antibody-containing supernatants were passed through 0.22µm filters. Antibodies were then purified using protein A affinity columns following the manufacturer’s protocol (Cytiva, Marlborough, MA, USA). Purified antibodies were dialyzed against PBS, run on SDS-PAGE and stained with Coomassie blue to assess for purity and integrity, before being aliquoted and stored at −80°C. The A32, 17b and 246D Fab fragments were generated by proteolytic digestion with immobilized papain (ThermoFisher Scientific) of 10mg/ml solutions of each antibody. Fab fragments were then purified with protein A followed by size exclusion chromatography (SEC) on a Superdex 200 increase 10/300 column (Cytiva) equilibrated in PBS.

### Plasma and antibodies

Plasma from PWH was obtained from the Canadian HIV and Aging Cohort Study (CHACS) [32] (Information on plasma donors can be found in Table **S3**). Plasma samples were collected, heat-inactivated for 1 h at 56°C and stored at −80°C until further use. Cell surface Env expression was measured using the 2G12 monoclonal antibody recognizing the gp120 outer domain. Cell surface CD4 expression was measured using a mouse monoclonal anti-CD4 antibody (clone OKT4, eBioscience, Catalog #14-0048-82) and cell surface BST-2 expression was measured using a mouse monoclonal anti-human CD317 antibody (clone RS38E, Catalog #348401). Alexa Fluor 647-conjugated anti-human IgG (Thermo Fisher Scientific, Cat #A-21445) was used as a secondary antibody to detect plasma binding and 2G12 binding, while the Alexa Fluor 647-conjugated anti-mouse IgG (Thermo Fisher Scientific, Cat # A-21235) was used as a secondary antibody to detect cell surface CD4 and BST-2 binding by flow cytometry. The PE-conjugated anti-HIV p24 antibody (clone KC57-RD1, Beckman Coulter, Cat # 6604667) was used to identify p24^+^ monocytes in the ADCP assay and was used in combination with the FITC-conjugated anti-human CD4 antibody (clone OKT4, Biolegend, Cat # 6604667) to identify productively-infected cells in cell-surface staining experiments as previously described [33].

### Small CD4-mimetic compound

Synthesis of the small CD4-mimetic compound CJF-III-288 has been previously described [20]. CJF-III-288 was dissolved to a stock concentration of 10mM in DMSO and stored at –80°C. The compound was then diluted to either 50µM (*ex vivo*-expanded CD4^+^ T cells), 10µM (HIV-1_AD8_, HIV-1_CH077T/F_, HIV-1_YU2_ and HIV-1_MM33T/F_) or 1µM (HIV-1_CH058T/F_) in PBS for cell surface staining experiments or in complete RPMI-1640 medium for ADCP experiments.

### Proviral constructs

Plasmids encoding the full length infectious molecular clones (IMCs) for HIV-1_AD8_ and HIV-1_YU2_ were previously described [12]. Plasmids encoding the IMCs of the HIV-1_CH058T/F_, HIV-1_CH077T/F_ and HIV-1_MM33T/F_ transmitted founder viruses and HIV-1_CH167_ were obtained from Dr Beatrice Hahn and described elsewhere [34–36]. The Vpu-deleted IMCs for HIV-1_CH058T/F_, HIV-1_CH077T/F_ and HIV-1_CH167_ were described in [37]. The Nef-deleted IMCs for HIV-1_CH058T/F_, HIV-1_CH077T/F_ and HIV-1_CH167_ were described in [38]. The Nef and Vpu-deleted IMCs for HIV-1_AD8_ and HIV-1_YU2_ were provided by Dr Frank Kirchhoff and described in [12]. The Nef and Vpu-deleted IMC for HIV-1_CH167_ was also provided by Dr Frank Kirchhoff. The Nef and Vpu-deleted IMCs for HIV-1_CH058T/F_ and HIV-1_CH077T/F_ were previously described in [38] and [39] respectively. The IMC encoding the Nef and Vpu-deleted HIV-1_CH058T/F_ harboring the Env D368R mutation was generated by site-directed mutagenesis by inserting the Env D368R mutation into the Nef and Vpu-deleted HIV-1_CH058T/F_ backbone using the QuikChange II XL site-directed mutagenesis protocol (Agilent, Cat# 200521). Sequences for all plasmids were verified by automated DNA sequencing.

### Viral production and infections

VSV-G-pseudotyped viral particles were produced by co-transfecting HEK293T cells with IMC-encoding plasmids and a VSV-G encoding plasmid at a ratio of 3:1 using polyethylenimine (PEI) in serum-free DMEM. Four hours post-transfection, media was changed for DMEM supplemented with 5% FBS and 100U/ml penicillin-streptomycin. Forty-eight hours post-transfection, supernatants were collected, cleared by low-speed centrifugation, and viral particles were concentrated by ultracentrifugation over a 20% sucrose cushion at 100,605g for 1h at 4°C. Viral pellets were resuspended in complete RPMI 1640 and stored at -80°C until further use. All viral productions were titrated on primary CD4^+^ T cells to obtain around 8-15% infected (p24^+^) cells after a forty-eight hours infection. For infection, activated primary CD4^+^ T cells were plated in a 96-well V-bottom plate at a concentration of 2 million cells/ml in complete RPMI 1640 medium supplemented with rIL-2 and appropriate viral titer. Cells were spinoculated at 800g for 1h and were incubated at 37°C for forty-eight hours.

### Interferon treatment

IFN-β (Rebif; EMD Serono Inc.) was reconstituted in RPMI-1640 at a concentration of 44µg/µl, aliquoted and stored at –80°C until further use. HIV-1-infected CD4^+^ T cells were treated with 2ng/µl of IFN-β twenty-four hours after infection, as previously reported [40]. This time represents twenty-four hours before cell surface staining and ADCP experiments.

### Cell-surface and intracellular staining

Forty-eight hours post-infection, infected primary CD4^+^ T cells were collected, washed with PBS and plated in 96-well V-bottom plates. For experiments with CJF-III-288, the appropriate concentration of the compound or equivalent volume of DMSO was added to the cells before plating. Cells were then incubated with either plasma (1:500 dilution), the 2G12 anti-Env monoclonal antibody (5µg/ml), an uncoupled anti-CD4 antibody (1µg/ml) or an uncoupled anti-human CD317 antibody (1µg/ml) for forty-five minutes at 37°C. Cells were washed twice with PBS and incubated at room temperature for twenty minutes with the appropriate secondary antibody mix: cells stained with plasma or 2G12 were incubated with Alexa Fluor 647-conjugated anti-human IgG (2µg/ml), LIVE/DEAD viability dye (1:1000 dilution, Thermo Fisher Scientific, Cat #L34957) and FITC-conjugated anti-CD4 (1:500 dilution). Cells stained with anti-CD4 or anti-CD317 were stained with Alexa Fluor 647-conjugated anti-mouse IgG (2µg/ml) and LIVE/DEAD viability dye (1:1000 dilution). After incubation, cells were washed twice with PBS, fixed/permeabilized in Cytofix/Cytoperm Fixation/Permeabilization Kit (BD Biosciences, Cat #554714) for at least twenty minutes and stained intracellularly with the PE-conjugated mouse anti-p24 monoclonal antibody (1:200 dilution). Samples were acquired on a Fortessa flow cytometer (BD Biosciences), and data analysis was performed using FlowJo v10.5.3. Plasma binding and 2G12 binding was determined by measuring the median fluorescence intensity of Alexa Fluor 647 from the productively infected living cells (LIVE/DEAD^low^, CD4^low^, p24^+^). CD4 and BST-2 levels at the surface of infected or non-infected cells were measured by the median fluorescence intensity of Alexa Fluor 647 in living p24^+^ or p24^-^ cells respectively.

### FACS-based ADCP assay

Primary CD4^+^ T cells were infected as described above. Sixteen hours before the ADCP assay, primary CD14^+^ CD16^-^ monocytes were isolated by negative selection, as described above, and rested overnight at 37°C at a concentration of 1 million cells/ml in RPMI 1640 supplemented with 20% FBS and 100U/ml penicillin-streptomycin. Forty-eight hours after infection, HIV-1-infected CD4^+^ T cells and rested primary monocytes were collected and counted. For experiments performed with CJF-III-288, productively infected p24^+^ CD4^low^ cells were enriched by depleting the CD4^high^ cells using the Dynabeads® CD4^+^ positive selection kit (Thermo Fisher Scientific, Cat #11145D) as previously described [11]. Infected CD4^+^ T cells were stained with the cell proliferation dye eFluor450 (Thermo Fisher Scientific, cat # 65-0842-90) diluted at 1:500 in PBS and used as target cells. Primary monocytes were stained with the cell proliferation dye eFluor670 (Thermo Fisher Scientific, cat #65-0840-90) diluted at 1:1000 in PBS and used as effector cells. Cells were stained for twenty minutes, washed twice with complete RPMI 1640 and resuspended in complete RPMI 1640. For experiments with CJF-III-288, the appropriate concentration of the compound or equivalent volume of DMSO was added to the cells before plating. For Fab blocking experiments, CJF-III-288-treated cells were incubated with A32 Fab (40µg/ml), 17b Fab (40 µg/ml), 246 Fab (80 µg/ml) or the combination of all Fab fragments for fifteen minutes at room temperature. Stained target cells were plated in 96-well V-bottom plates and incubated with plasma (1:500 dilution) for forty-five minutes at 37°C. After incubation, target cells were washed twice with complete RPMI 1640 to remove unbound antibodies. Antibody-opsonized target cells were subsequently put in coculture with the effector cells in 96-well U-bottom plates at a 3:1 target to effector ratio for two hours at 37°C. After incubation, cells were washed with PBS, fixed/permeabilized in Cytofix/Cytoperm Fixation/Permeabilization Kit and stained intracellularly using the PE-conjugated mouse anti-p24 antibody. Samples were acquired on a Fortessa cytometer and data analysis was performed using FlowJo v10.5.3. The ADCP index was calculated as follows: (% of PE^+^ effectors in targets + effectors + plasma) / (% of PE^+^ effectors in targets + effectors).

### Image-capture flow cytometry

Primary CD4^+^ T cells were infected with Nef and Vpu-deleted HIV-1_CH058T/F_. Forty-eight hours post-infection, HIV-1-infected CD4^+^ T cells and primary monocytes, which were isolated and maintained as described above, were collected and counted. CD4^+^ T cells were stained with the cell proliferation dye eFluor450 in PBS at a 1:1000 dilution and monocytes were stained with the cell proliferation dye CFSE (Invitrogen, Cat#C34554) at a 1:8000 dilution in PBS. Cells were stained for twenty minutes, washed twice with complete RPMI 1640 and resuspended in complete RPMI 1640. CD4^+^ T cells were plated in 96-well V-bottom plates and incubated with plasma (1:500 dilution) for forty-five minutes at 37°C. After incubation, cells were washed twice with complete RPMI 1640 to remove unbound antibodies and were put in coculture with CFSE-stained monocytes in 96-well U-bottom plates at a 3:1 target to effector ratio for two hours at 37°C. After incubation, cells were washed with PBS, fixed/permeabilized in Cytofix/Cytoperm Fixation/Permeabilization Kit and stained intracellularly using the PE-conjugated mouse anti-p24 antibody at a 1:200 dilution. After staining, samples were resuspended in PBS and transferred to 1.5ml microcentrifuge tubes. Samples were run on an ImageStreamX MKII flow cytometer and data analysis was performed with the IDEAS software (Amnis).

### Quantification and statistical analysis

Statistical analyses were performed using GraphPad Prism version 10.2.0. Each data set was tested for statistical normality to determine the appropriate statistical test to use (parametric or nonparametric). Statistical details of experiments are indicated in the figure legends. p values < 0.05 were considered significant; significance values are indicated as *p < 0.05, **p < 0.01, ***p < 0.001, ****p < 0.0001 and ns: nonsignificant.

## RESULTS

### Accessory proteins Nef and Vpu protect HIV-1-infected cells from ADCP

It is well established that accessory proteins Nef and Vpu allow to escape ADCC responses mediated by PWH plasma, notably through the downregulation of CD4, but also due to the ability of Vpu to downregulate BST-2 (Tetherin, CD317) at the surface of infected cells [6, 7, 12, 31, 41, 42]. Therefore, we wanted to determine if similar mechanisms were involved in the protection of infected cells from ADCP-mediated killing. To answer this, we developed a FACS-based assay to accurately measure ADCP-mediated killing of HIV-1-infected cells (**Figure 1A**). We started by infecting primary CD4^+^ T cells with primary HIV-1 isolates for forty-eight hours. These cells, which served as target cells in our assay, were stained with the eFluor450 cell viability dye and incubated with plasma from PWH to allow antibody opsonization. Plasma from people without HIV-1 (PWoH) was used as a negative control. Primary monocytes were isolated by negative selection (see Material and Methods) from autologous PBMCs, stained with the eFluor670 cell proliferation dye and served as effector cells. Purity of isolated primary monocytes was confirmed by flow cytometry (**Figure S1**). Opsonized target cells and effector cells were put in co-culture for two hours at a ratio of 3:1 (target to effector) and were subsequently stained intracellularly for p24. ADCP activity represents the ratio of p24^+^ monocytes in conditions with targets + effectors + plasma over the p24^+^ monocytes in targets + effectors alone (see gating strategy in **Figure S2**). To assess if Nef and Vpu could protect infected cells from ADCP mediated by PWH plasma, we infected primary CD4^+^ T cells with either wild-type (WT) or Nef and Vpu-deleted (Nef-Vpu-) HIV-1 isolates and evaluated how these two accessory proteins impacted recognition and ADCP-mediated killing. For all four primary isolates tested, we observed that cells infected with Nef-Vpu- viruses were significantly more recognized by plasma from PWH (see supplemental tables for participants demographics) compared to WT-infected cells (**Figure 1B**), in line with previously reported data [7]. This is due to an increase in CD4 expression at the surface of infected cells (**Figure 1E**), which enables Env to sample more “open” conformations recognized by nnAbs present in PWH plasma, but also due to the increase in cell surface BST-2 expression (**Figure 1F**), which traps newly budded viral particles and results in the accumulation of Env at the cell surface. Cells infected with WT viruses were not readily eliminated by ADCP in presence of PWH plasma. Interestingly, deletion of Nef and Vpu rendered infected cells significantly more susceptible to ADCP mediated by PWH plasma (mean ADCP index for all viruses: 2.286), suggesting a role for these accessory proteins in the protection of infected cells from this Fc-effector function (**Figure 1C**). As expected, plasma binding strongly correlated with ADCP activity (**Figure 1D**), suggesting that increased recognition of infected cells by antibodies present in plasma renders them susceptible to ADCP-mediated killing.

**Figure 1.**
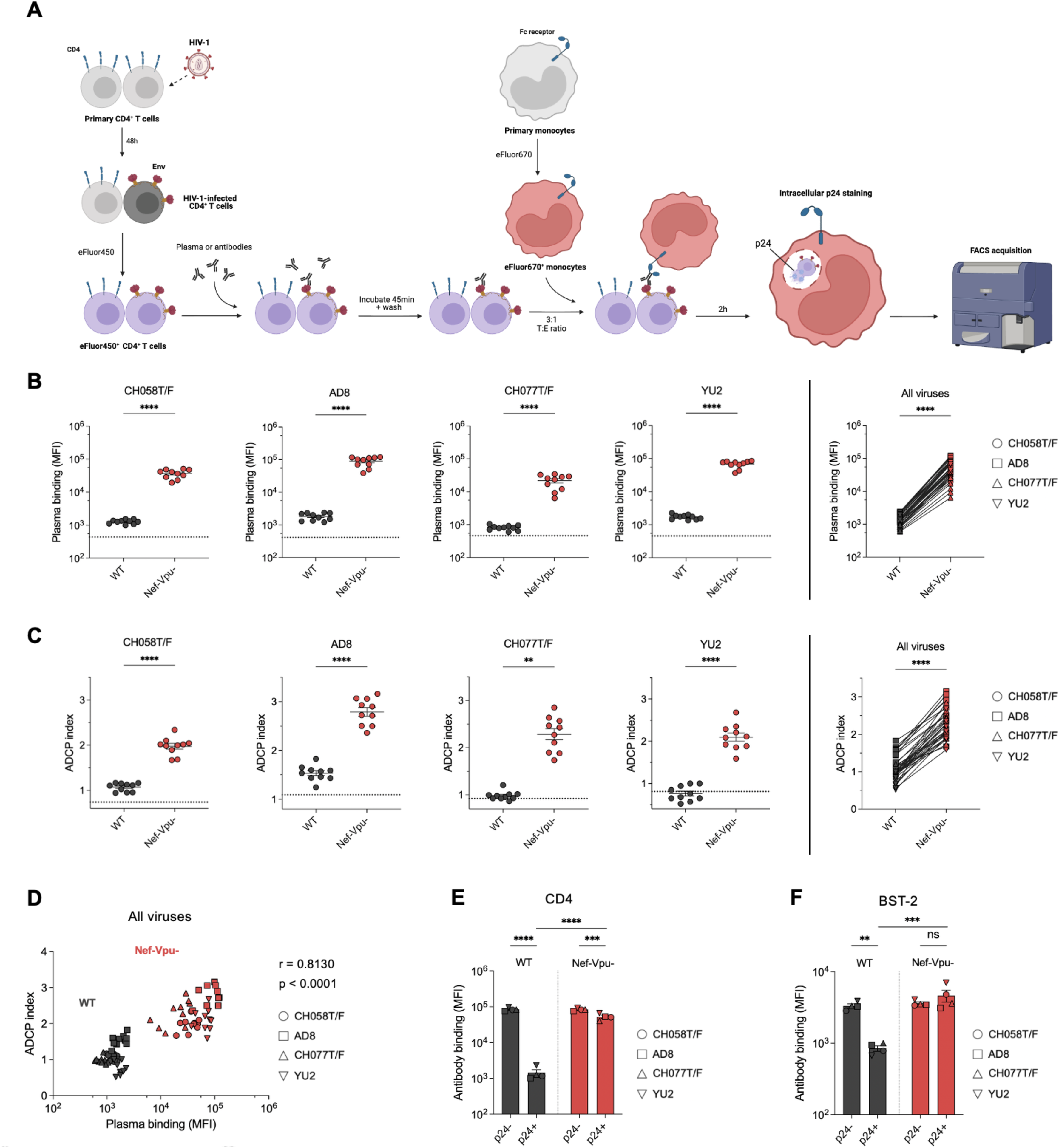
Nef and Vpu protect HIV-1-infected cells from ADCP mediated by PWH plasma. (**A**) Schematic representation of the FACS-based ADCP assay. Primary CD4^+^ T cells were infected with various HIV-1 isolates for forty-eight hours and used as target cells. On the day of the assay, HIV-1-infected CD4^+^ T cells were stained with the eFluor450 proliferation dye and incubated for forty-five minutes with plasma from PWH or PWoH. Autologous primary monocytes were used as effector cells and were stained with the eFluor670 proliferation dye. Monocytes were put in coculture with opsonized CD4^+^ T cells at a 3:1 target (T) to effector (E) ratio for two hours. After coculture, cells were fixed and stained intracellularly for the p24 antigen, and data was acquired by FACS. (**B**) Recognition of primary CD4^+^ T cells infected with wild-type (WT) or Nef and Vpu-deleted (Nef-Vpu-) HIV-1 isolates by plasma from 10 PWH. Dotted lines represent the mean plasma binding from 5 PWoH. (**C**) ADCP-mediated elimination of primary CD4^+^ T cells infected with WT or Nef-Vpu- HIV-1 isolates in presence of PWH plasma. ADCP index was calculated as the percentage of p24^+^ monocytes T + E + plasma samples divided by the percentage of p24^+^ monocytes in T + E alone. Dotted lines represent the mean ADCP index measured in presence of plasma from 5 PWoH. (**D**) Spearman correlation between PWH plasma binding and ADCP activity measured for all viruses. (**E-F**) Expression of CD4 (**E**) and BST-2 (**F**) at the surface of cells infected with WT or Nef-Vpu- HIV-1 isolates. Each data set was tested for normality, and statistical significance was determined by paired t-tests or Wilcoxon matched-pairs signed rank test for normally distributed data or non-normally distributed data respectively for B and C. Statistical significance was determined by two-way ANOVA for E and F (n=2).

We next visualized the ADCP events by image capture flow cytometry. Briefly, primary CD4^+^ T cells infected with Nef and Vpu-deleted HIV-1_CH058T/F_, were stained with eFluor450 and cocultured with CFSE-stained monocytes in presence of PWH plasma for two hours. Cells were subsequently stained intracellularly with a PE-coupled anti-p24 antibody and run on a ImageStream MKII flow cytometer. As expected, when infected cells and monocytes were co-cultured in the presence of PWH plasma, we observed monocytes that were positive for the CD4^+^ T cell marker and that also contained intracellular p24^+^ puncta (**Figure 2**), representing monocytes that had phagocytosed HIV-1-infected cells. Overall, here we show that Nef and Vpu protect HIV-1-infected cells from ADCP.

**Figure 2.**
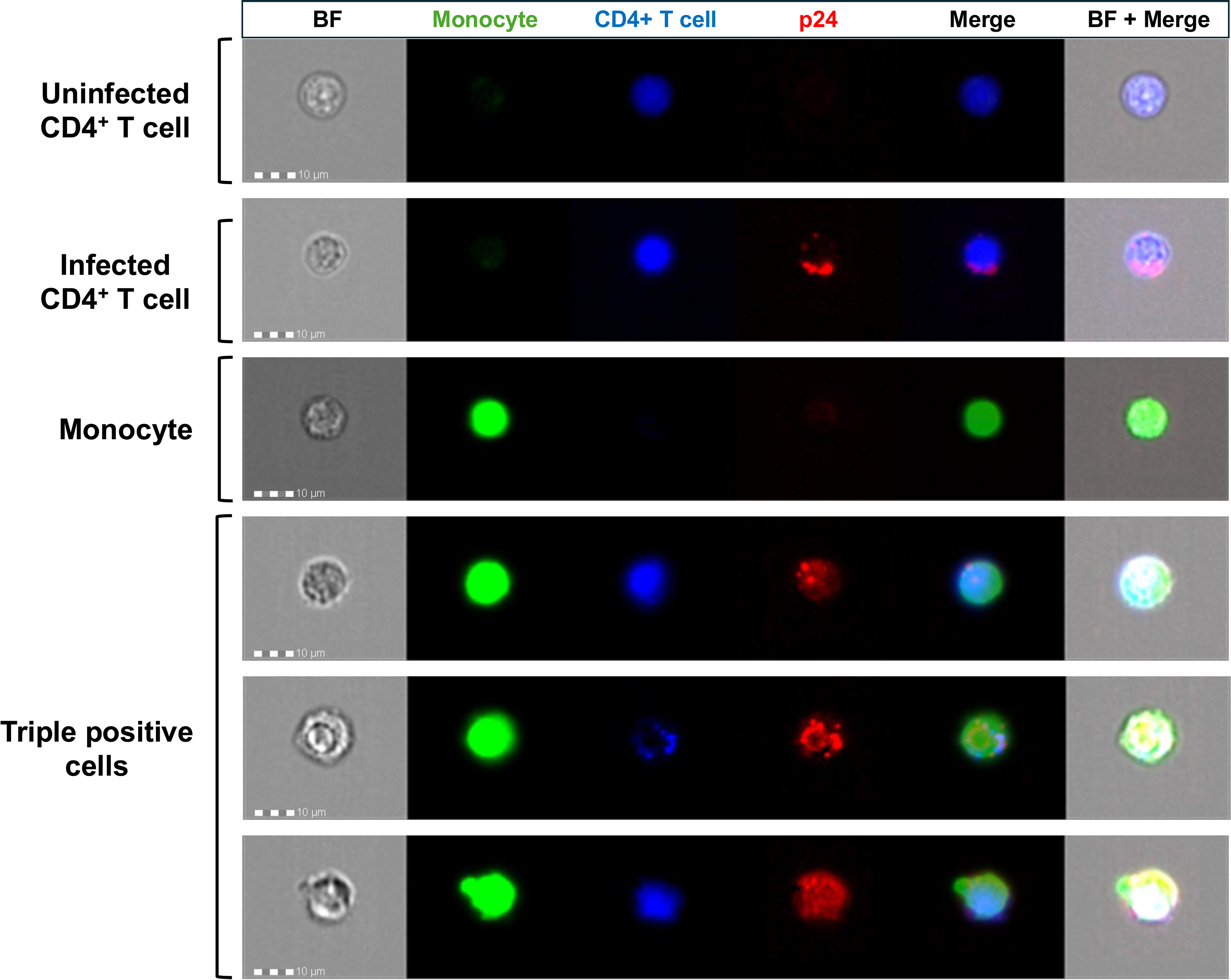
Image capture flow cytometry images of phagocytosis of HIV-1-infected CD4^+^ T cells by monocytes. Primary CD4^+^ T cells were infected with Nef-Vpu- HIV-1_CH058T/F_ and were labelled with the cell proliferation dye eFluor450 (blue). Primary monocytes were labelled with the cell proliferation dye CFSE (green) and both cell types were put in coculture in presence of PWH plasma for two hours. Cells were then fixed and stained intracellularly for the p24 antigen (red). Shown are images of primary monocytes alone, CD4^+^ T cells alone or cocultured cells in presence of PWH plasma. Cells were selected for focus, size and aspect ratio for phenotypic analysis. BF: brightfield.

### Env conformation modulates the susceptibility of HIV-1-infected cells to ADCP

Next, we sought to dissect the individual roles of Nef and Vpu in protecting HIV-1-infected cells from ADCP mediated by plasma from PWH. To do so, we infected CD4^+^ T cells with WT HIV-1_CH058T/F_ or with viruses deleted for either Nef (Nef-), Vpu (Vpu-) or both accessory proteins (Nef-Vpu-). Deletion of a single accessory protein increased recognition of infected cells by plasma from PWH (**Figure 3A**). For cells infected with the Nef- virus, this was likely explained by an increase in cell surface expression of CD4, allowing Env to sample more “open” conformations, thereby exposing epitopes recognized by CD4i Abs which are abundant in the plasma of PWH. Deletion of Vpu enhanced the levels of both CD4 and BST-2. Similar to Nef deletion, the increase in CD4 levels leads to the exposure of CD4i epitopes while BST-2 upregulation results in the accumulation of Env at the cell surface, due to trapped viral particles (**Figure 3C-D-E**). Surprisingly, although deletion of Nef or Vpu led to similar increases in plasma binding, deletion of Vpu caused a significantly greater increase in ADCP activity, which reached similar levels than those observed with the Nef-Vpu-virus (**Figure 3B**). These results suggest that Vpu has a greater contribution than Nef in protecting cells from ADCP. To confirm that this phenotype was not unique to HIV-1_CH058T/F_ we repeated the experiment with two additional HIV-1 isolates (CH077T/F and CH167). Although Nef also contributed to protect infected cells from ADCP, the effect of Vpu was significantly higher (**Figure S3**). To ascertain the role of Env-CD4 interaction in ADCP, we introduced the Env D368R mutation, which abrogates Env-CD4 interaction [43], in the Nef-Vpu- HIV-1_CH058T/F_ backbone. As previously reported [6, 7, 12], introduction of the D368R mutation slightly increased Env levels at the surface of Nef-Vpu- infected cells (**Figure 3E**). In agreement with the critical role that the D368 residue of Env plays in CD4 interaction, cells infected with the Nef-Vpu- D368R virus were not efficiently recognized by PWH plasma due to the inability of Env to interact with CD4, thereby preventing the exposure of CD4i epitopes (**Figure 3A**). Despite poor recognition of these cells, they remained susceptible to ADCP to similar levels than Vpu- infected cells (**Figure 3B**), thus suggesting that additional mechanisms, likely a cell host protein targeted by Vpu, contribute to the protection of HIV-1-infected cells from ADCP.

**Figure 3.**
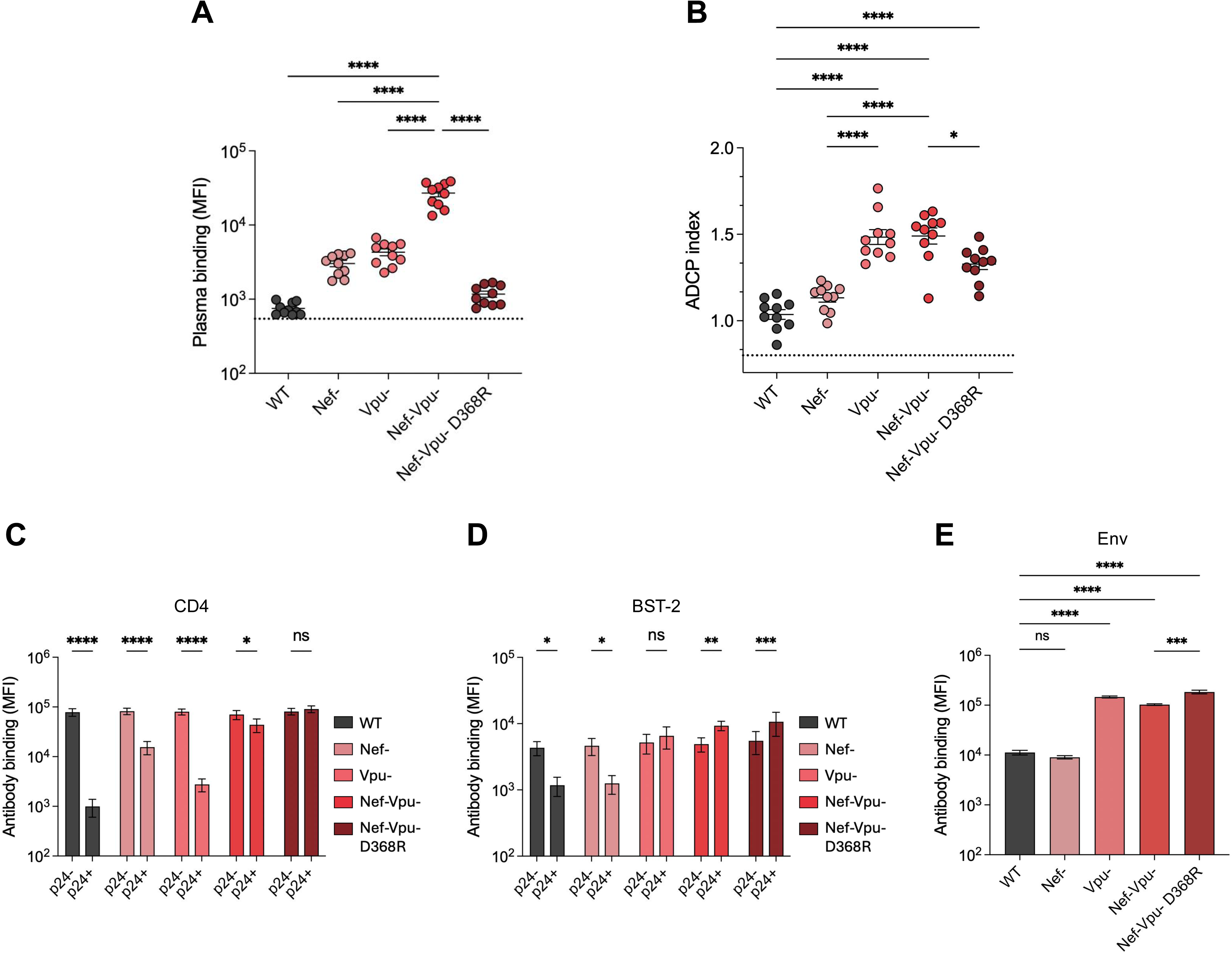
Env-CD4 interaction facilitates recognition and ADCP-mediated killing of infected cells by PWH plasma. (**A**) Recognition and (**B**) PWH plasma-mediated ADCP killing of CD4^+^ T cells infected with wild-type (WT) HIV-1_CH058T/F_, a virus deleted for either Nef (Nef-), Vpu (Vpu-), both accessory proteins (Nef-Vpu-) or deleted for Nef and Vpu in addition to harboring the Env D368R mutation (Nef-Vpu- D368R). Dotted lines represent plasma binding or ADCP index that was measured with plasma from PWoH. (**C-D**) Expression of CD4 (**C**) and BST-2 (**D**) at the surface of p24^-^ or p24^+^ cells infected with the indicated virus. (**E**) Cell surface expression of Env measured by binding of the 2G12 monoclonal antibody. Statistical significance was determined by one-way ANOVA for A-B-E and two-way ANOVA for C-D (n=3).

### “Opening up” Env with CD4mc sensitizes cells to ADCP

Our experiments with the D368R mutant support a major role for Env conformation in ADCP susceptibility. To confirm this, we used a potent indoline CD4mc, CJF-III-288, which was recently shown to sensitize HIV-1-infected cells to ADCC mediated both by PWH plasma and various combinations of nnAbs *in vitro* [17–20]. Briefly, we infected primary CD4^+^ T cells with five different primary HIV-1 isolates and evaluated if CJF-III-288 sensitized these cells to ADCP mediated by plasma from PWH. CJF-III-288 significantly enhanced recognition cells and elimination of infected cells by ADCP for all the viruses tested (**Figure 4A-D**). As expected, we noted a significant positive correlation between plasma binding and ADCP activity (**Figure 4E**), thus confirming the central role of Env conformation on the susceptibility of HIV-1-infected cells to ADCP.

**Figure 4.**
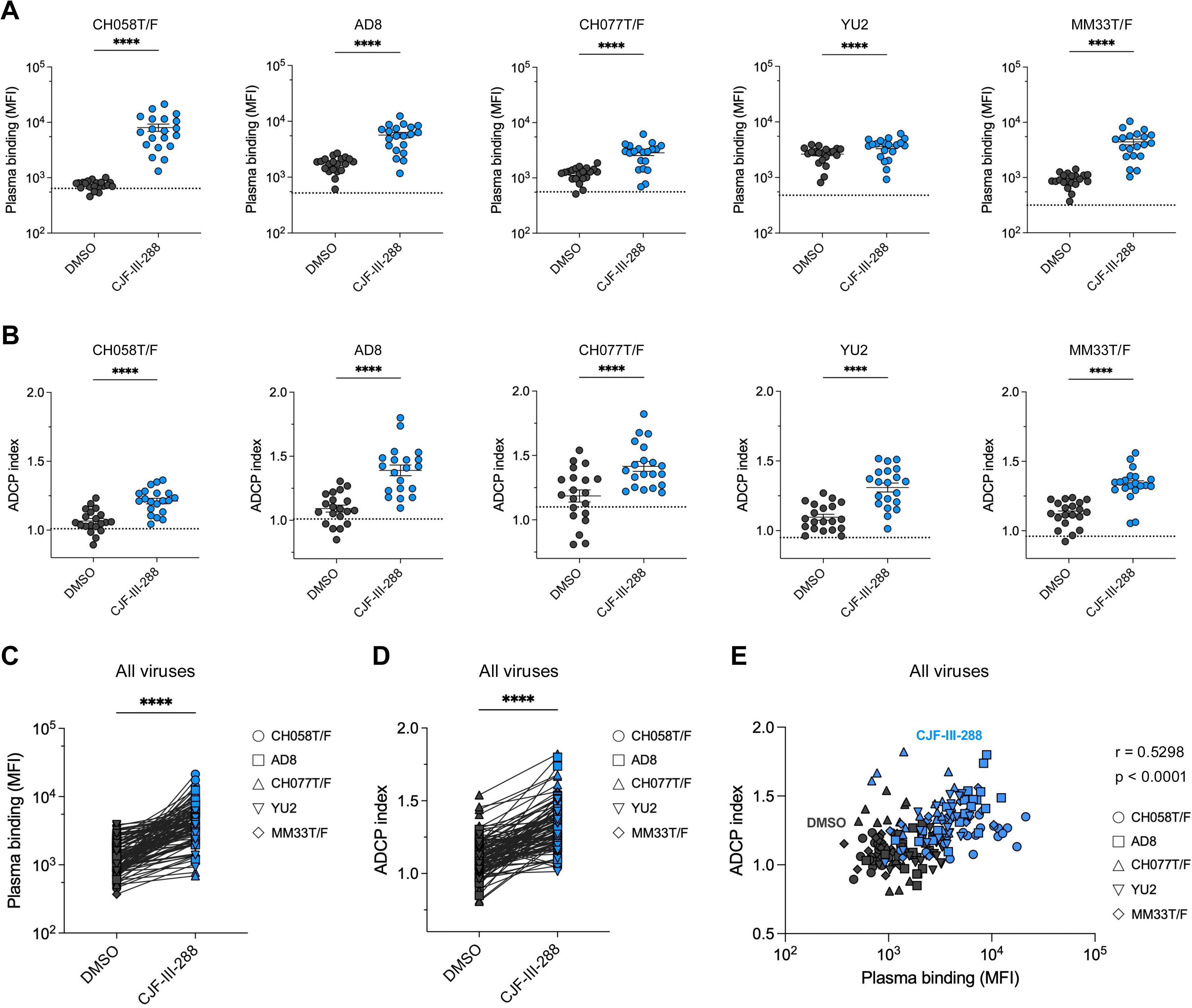
CJF-III-288 sensitizes HIV-1-infected cells to ADCP mediated by PWH plasma. (**A**) Recognition and (**B**) ADCP-mediated elimination of HIV-1-infected CD4^+^ T cells by PWH in presence of the CD4-mimetic compound CJF-III-288 (1µM for HIV-1_CH058T/F_, 10µM for HIV-1_AD8_, HIV-1_CH077T/F_, HIV-1_YU2_ and HIV-1_MM33T/F_) or equivalent volume of vehicle (DMSO). Dotted lines represent the plasma binding or ADCP activity that was measured with plasma from PWoH. (**C-D**) Compilation of recognition (**C**) and ADCP activity (**D**) mediated by PWH plasma for all the tested HIV-1 isolates. (**E**) Spearman correlation between PWH plasma binding and ADCP activity measured for all viruses. Each data set was tested for normality and statistical significance was determined by paired t-tests or Wilcoxon matched-pairs signed rank test for normally distributed data or non-normally distributed data respectively (n=2).

Next, we investigated whether we could translate these findings to *ex vivo*-expanded CD4^+^ T cells from PWH under ART. To do so, we isolated CD4^+^ T cells from the PBMCs of five PWH under ART (information on participants can be found in **Table S2**), activated them with PHA-L and cultured them in presence of IL-2, monitoring p24 levels to assess viral replication. Cells were cultured until we detected at least 5% of p24^+^ cells (**Figure S4**), upon which we performed our assays. Treatment of the *ex vivo*-expanded CD4^+^ T cells with CJF-III-288 led to a significant increase in binding by heterologous and autologous plasma (autologous plasma for participants 4 and 5 were not available), which is in line with our data using *in vitro*-infected cells (**Figure 5A-C**). Importantly, treatment with the CD4mc significantly increased ADCP-mediated killing of infected cells in presence of heterologous plasma for Participants 1, 3, 4 and 5 (**Figure 5B**). Although the increase in ADCP activity was not statistically significant for Participant 2, we still observed increased ADCP in presence of the participant’s autologous plasma, and when combining the data from all the participants together, we observed a significant increase in ADCP-mediated killing in presence of CJF-III-288 (**Figure 5D**). Once again, plasma binding positively correlated with ADCP activity (**Figure 5E**). Altogether, these results demonstrate that CD4mc such as CJF-III-288 can be used to sensitize both *in vitro*-infected and *ex vivo*-expanded CD4^+^ T cells from PWH under ART to ADCP mediated by plasma from PWH.

**Figure 5.**
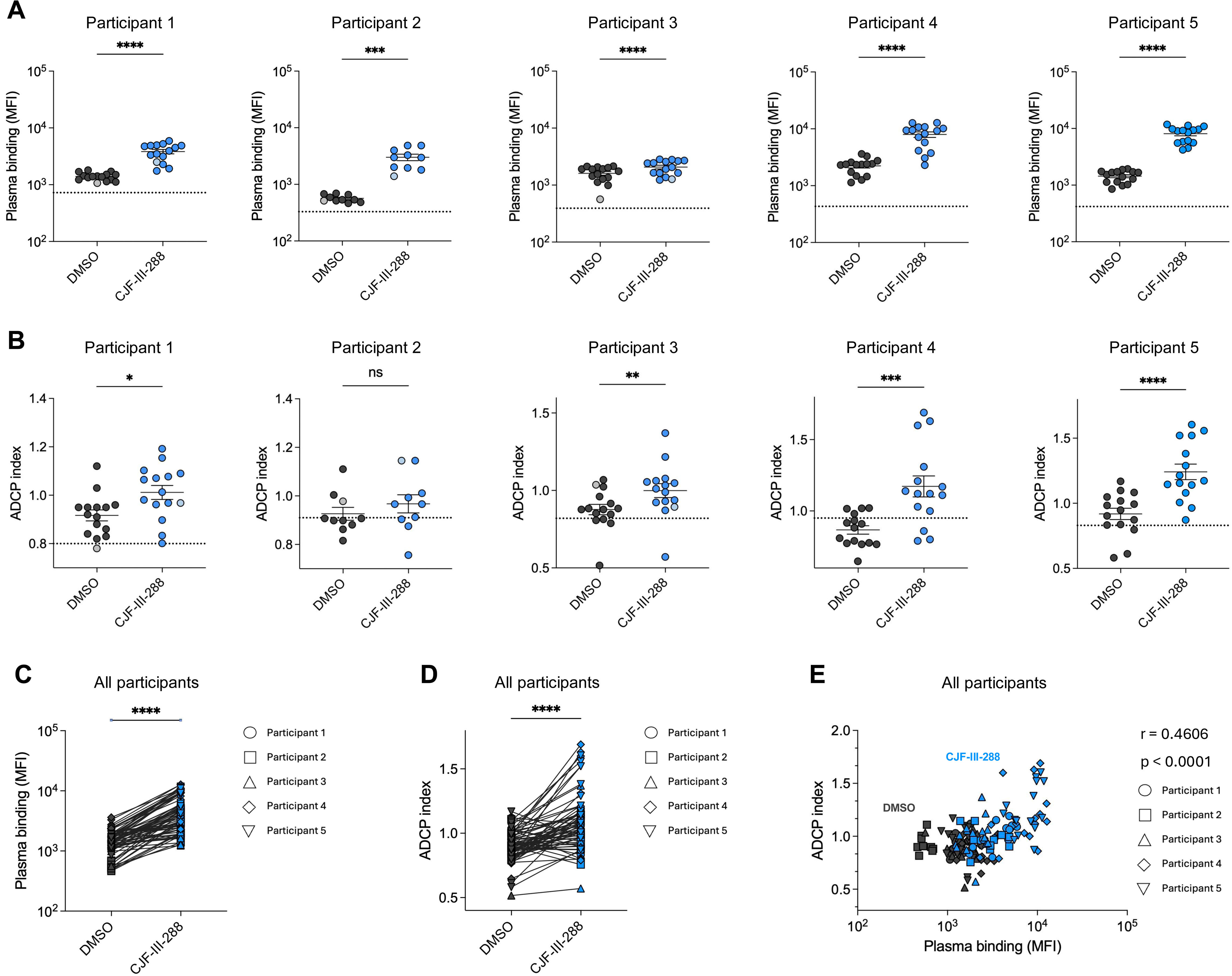
CJF-III-288 sensitizes *ex vivo*-expanded CD4^+^ T cells from PWH under ART to ADCP mediated by PWH plasma. (**A**) PWH plasma-mediated recognition and (**B**) ADCP killing of *ex vivo*-expanded CD4^+^ T cells from four PWH under ART in presence of DMSO or the CD4mc CJF-III-288 (50µM). Data points in lighter colors represent autologous plasma binding and ADCP activity in presence of DMSO (light grey) or CJF-III-288 (light blue), except for participants 4 and 5, for which the autologous plasma was not available. (**C-D**) Compilation of plasma binding (**C**) and ADCP activity (**D**) mediated by PWH plasma on ex vivo-expanded CD4^+^ T cells from PWH under ART for all participants. (**E**) Spearman correlation between PWH plasma binding and ADCP activity measured for all participants. Each data set was tested for normality and statistical significance was determined by paired t-tests or Wilcoxon matched-pairs signed rank test for normally distributed data or non-normally distributed data respectively.

### Anti-cluster A antibodies play an important role in ADCP mediated by plasma from PWH

The composition of plasma from PWH is heterogenous and the composition of anti-Env antibodies mainly includes various families of CD4i antibodies such as anti-cluster A, anti-coreceptor binding site (CoRBS) and anti-gp41 cluster I antibodies [18]. We previously evaluated the ability of plasma from 50 PWH to mediate ADCC in presence of CJF-III-288 and observed that although all of these antibody families played a role in plasma-mediated ADCC, their contribution varied [18]. Here, we evaluated how each of these families of antibodies was involved in ADCP-mediated killing of infected cells by performing a Fab competition experiment. Briefly, HIV-1-infected CD4^+^ T cells were incubated with CJF-III-288 and either A32 Fab (cluster A), 17b Fab (CoRBS), 246D Fab (gp41 cluster I) or the combination of all three specificities. These Fab fragments bind to their respective epitopes upon Env opening with the CD4mc and thereby “block” antibodies in the plasma with the same specificity from binding Env (**Figure 6A**). The reduction of the ADCP response in presence of these Fab fragments represents the relative importance of each family of antibodies to ADCP. We observed that pre-incubation of infected cells with A32 Fab fragments significantly decreased the ability of plasma to mediate ADCP in presence of CJF-III-288 (**Figure 6B**). Incubation with 17b or 246D Fab fragments also decreased the ADCP response on their own, albeit to a lesser extent than A32. The combination of all three Fab fragments further decreased the ADCP response mediated by PWH plasma in presence of the CD4mc, as was observed for incubation with the A32 Fab fragment alone. These results show that all three of these families of CD4i antibodies contribute to the level of ADCP response mediated by PWH plasma, but that this Fc-effector function is mostly driven by anti-cluster A antibodies.

**Figure 6.**
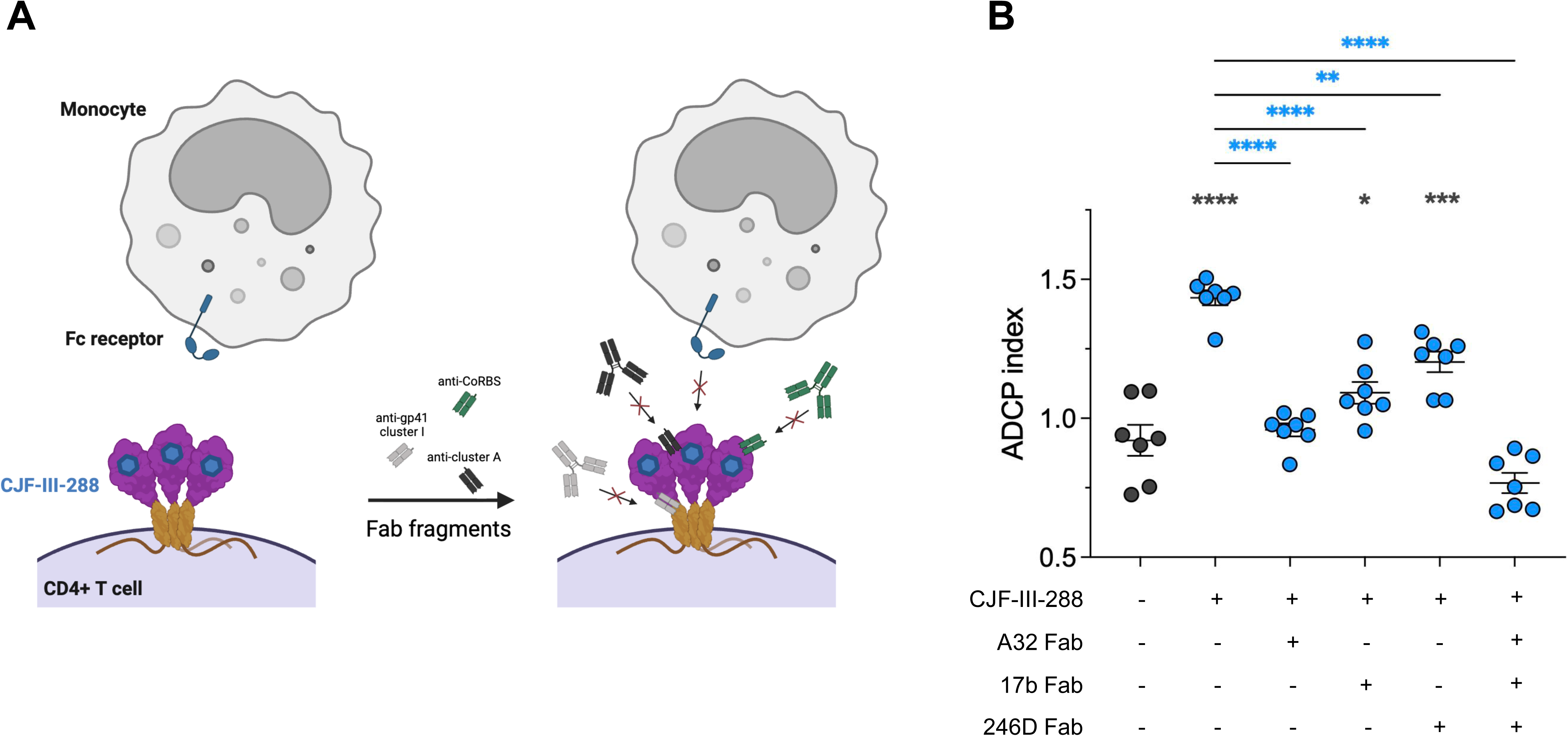
Anti-cluster A antibodies play a major role in PWH plasma mediated ADCP. (**A**) Schematic representation of Fab fragment blocking of HIV-1-infected cells. Pre-treatment of HIV-1-infected CD4^+^ T cells with CJF-III-288 and Fab fragments allows them to bind to their respective epitopes and prevents antibodies of the same class in plasma from binding to their epitopes. This also prevents Fc receptor-mediated recruitment of effector cells and thus the elimination of infected cells by ADCP. (**B**) ADCP-mediated elimination of HIV-1_CH058T/F_-infected CD4^+^ T cells by PWH plasma in presence of CJF-III-288 (1µM) with or without pre-incubation with A32 Fab (anti-cluster A), 17b Fab (anti-CoRBS) or 246D Fab (anti-gp41 cluster I). Statistical significance was determined by one-way ANOVA with Tukey’s multiple comparisons test. Statistics in blue represent comparisons with the CJF-III-288 without Fab condition and statistics in grey represent comparisons with the DMSO condition (n=3).

### IFN-**𝛃** enhances ADCP activity of PWH plasma in presence of CJF-III-288

Thus far, we have shown CJF-III-288 could sensitize infected cells to ADCP, although the magnitude of the response observed was lower than what was observed for cells infected with Nef-Vpu- viruses. We therefore wondered if we could employ additional strategies to increase the susceptibility of WT-infected cells to this Fc-effector function. BST-2 antagonism by Vpu has been shown to protect HIV-1-infected cells from ADCC mediated both by bNAbs [42] and by sera from PWH in presence of CD4mc [31]. In both experimental settings, treatment of infected cells with type I interferon (IFN), which is known to induce BST-2 expression [31, 41], increased their susceptibility to ADCC-mediated killing. To assess if a similar strategy could be used to enhance ADCP responses in presence of the CD4mc, we treated CD4^+^ T cells infected with HIV-1_CH058T/F_ with IFN-β for 24 hours. As expected, IFN-β treatment significantly increased BST-2 expression at the surface of both infected and uninfected cells and also increased cell surface Env (**Figure S5**). The combination of CJF-III-288 and IFN-β significantly increased both recognition and ADCP-mediated killing of infected cells, both of which significantly correlated (**Figure 7**). Taken together, these results support that inducing BST-2 expression with IFN-β can lead to the accumulation of Env at the cell surface, thereby increasing the recognition and ADCP-mediated killing of infected cells by PWH plasma in presence of CJF-III-288.

**Figure 7.**
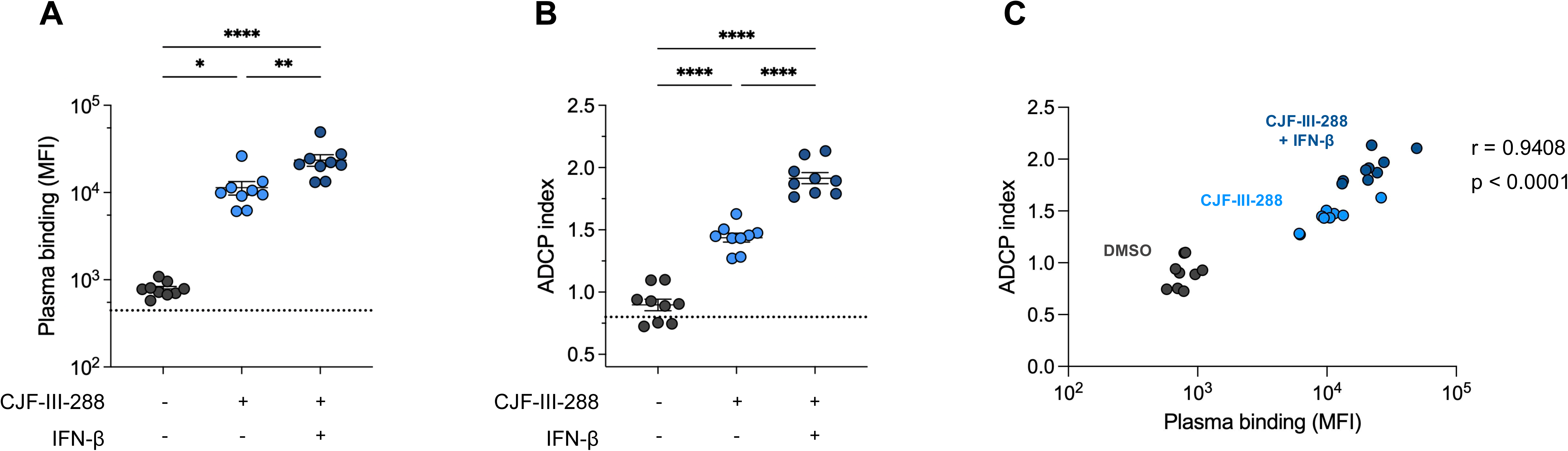
**IFN-β enhances the susceptibility of HIV-1-infected cells to ADCP mediated by plasma from PWH in presence of CJF-III-288**. (**A**) Recognition and (**B**) ADCP-mediated elimination of HIV-1_CH058T/F_-infected CD4^+^ T cells by plasma from PWH with or without treatment with CJF-III-288 (1µM) and IFN-β (2ng/ml). Dotted lines represent the plasma binding or ADCP activity that was measured with plasma from PWoH. Statistical significance was determined by one-way ANOVA with Tukey’s multiple comparisons test. (**C**) Spearman correlation between PWH plasma binding and ADCP activity with or without CJF-III-288 and IFN-β treatment (n=3).

## DISCUSSION

Although antiretroviral therapy suppresses viremia in most PWH [44], it does not decrease the size of the viral reservoir [45, 46]. A growing body of evidence highlights the potential of ADCC to target the reservoir, but the role of other Fc-effector functions such as ADCP remains less characterized. Previous studies analyzing immune correlates of protection among participants of HIV-1 vaccine trials as well as vaccine candidates in non-human primate challenge studies determined that elicitation of Env specific ADCP-mediating antibodies was associated with decreased risk of HIV-1 acquisition [28, 30] and that elicitation of anti-Env antibodies with high affinity binding to CD32 was associated to protection in the HVTN 505 trial [29]. However, the methods used in these studies to reach these conclusions did not directly assessed the ability of antibodies to induce the phagocytosis of HIV-1 infected cells, but rather measured the phagocytosis of beads coated with recombinant Env proteins, which present epitopes that differ from those found on membrane-bound Env trimers [25, 47].

In this work, we developed a FACS-based assay to measure the elimination of primary CD4^+^ T cells infected with primary HIV-1 isolates by ADCP, mediated by autologous monocytes. We observed that Nef and Vpu contribute to protect HIV-1-infected cells against ADCP mediated by antibodies present in the plasma of PWH, similar to what was reported for ADCC [6, 7, 12]. Indeed, we saw that cells infected with primary HIV-1 isolates were not readily phagocytosed in presence of PWH plasma, but that deletion of Nef and Vpu rendered them highly vulnerable to ADCP-mediated killing (**Figure 1**). This was in part linked to the ability of these accessory proteins to downregulate CD4, as cells infected with a Nef-Vpu- virus harboring the Env D368R mutation were less efficiently eliminated (**Figure 3**). This further emphasizes the central role that CD4 downregulation plays in promoting viral pathogenesis. Of note, our results also suggest that Vpu could protect infected cells from ADCP through additional mechanisms independent from CD4 downregulation. It is known that the Vpu-mediated downregulation of NK cell ligands such as NTB-A, PVR and CD48 at the surface of infected cells serves to protect infected cells from ADCC [40, 48–50]. It is therefore possible that Vpu could be involved in the regulation of other immunomodulating ligands regulating ADCP responses, although this remains to be explored further.

CD4mc induce conformational changes in the Env trimer similar to those that occur upon CD4 binding [19, 51, 52]. This stabilizes Env in more “open” downstream conformations and enables CD4i nnAbs to sequentially bind to the trimer and sensitize infected cells to elimination by ADCC [16, 53]. Here, we show that the indoline CJF-III-288 CD4mc sensitizes cells infected with five different primary HIV-1 isolates to ADCP-mediated killing by PWH plasma (**Figure 4**) and that this activity can be further enhanced by treating infected cells with IFN-β (**Figure 7**). Importantly, there have been reports that HIV-1 infection causes dysregulation in monocyte biology, notably by inducing their chronic activation, altering their cytokine profiles, and reducing their phagocytic capacities [54, 55]. Still, we observed that CJF-III-288 could sensitize *ex vivo*-expanded CD4^+^ T cells from PWH under ART to ADCP mediated by their autologous monocytes (**Figure 5**). This suggests that ADCP-based immunotherapies could be considered to eliminate infected cells in PWH despite reports of immune dysfunction affecting monocytes of PWH.

By performing Fab blocking experiments, we found that different families of antibodies present in PWH plasma had different contributions to ADCP activity, with anti-cluster A antibodies being the strongest inducers of this response (**Figure 6**). We previously studied the importance of various families of nnAbs present in PWH plasma in ADCC-mediated killing of HIV-1 infected cells [18]. Interestingly, we observed that CoRBS antibodies had a greater role in mediating this Fc-effector function, highlighting the fact that different classes of anti-Env antibodies can have different propensities to induce ADCC and ADCP. Whether this is due to a preferential angle of approach required for the Fc portion of antibodies to interact with CD16 versus CD32, or to steric hindrance related to the type and size of the effector cells primarily involved in these processes, remains to be elucidated.

ADCC and ADCP could both play complementary roles in the elimination of HIV-1 infected cells. For example, lymphoid tissues, which represent major anatomical sites harboring HIV-1 reservoir cells [56, 57], are mostly devoid of ADCC-mediating CD56^dim^ CD16^+^ NK cells [58, 59]. In contrast, monocytes are vastly abundant in lymph nodes and could serve as key players in eliminating HIV-1-infected cells in this compartment. As lymphoid tissues constitute major sites of viral rebound in SIV-infected macaques upon ART interruption [60], monocyte-mediated Fc-effector functions in these organs could be all the more important to achieve post-treatment HIV-1 control. Still, the ability of combinatorial nnAbs/CD4mc treatments to induce ADCP of HIV-1 infected cells *in vivo* remains to be thoroughly evaluated.

Overall, we show here that Nef and Vpu accessory proteins protect HIV-1-infected cells from ADCP. CD4mc-based strategies can be used to “open-up” Env, thereby harnessing the full potential of this Fc-effector function. By increasing the levels of Env available for CD4mc engagement, type I IFN can further enhance ADCP-mediated elimination of HIV-1 infected cells (**Figure 8**). Our findings warrant additional work to better understand the contribution of different Fc-effector functions in the elimination of HIV-1-infected cells, which could inform the development of new immunotherapies toward an HIV-1 cure.

**Figure 8.**
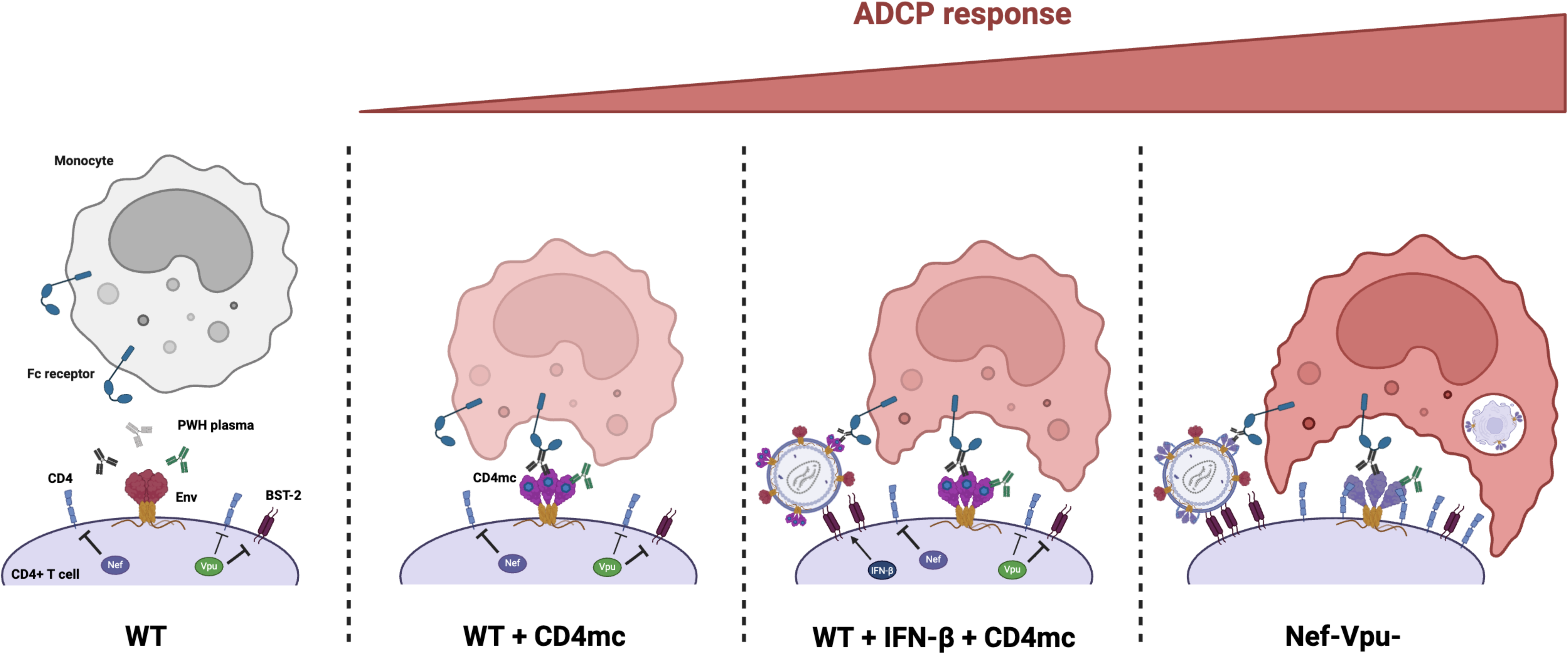
**Schematic of HIV-1-infected cells susceptibility to ADCP responses**. Cells infected with WT HIV-1 isolates are not efficiently recognized by plasma from PWH due to the protective effect of the accessory proteins Nef and Vpu. Utilizing CD4mc such as CJF-III-288 can force Env “opening”, thereby allowing PWH plasma to recognize infected cells and sensitize them to ADCP. The combination of CJF-III-288 with treatment of infected cells with IFN-β can further increase the ADCP activity of PWH plasma due to the induction of BST-2 expression, which increases Env expression at the surface of infected cells. The highest ADCP activity is achieved against cells infected with a Nef and Vpu-deleted virus, as these cells express large quantities of Env in its “open” conformation, leading to the highest recognition by PWH plasma.

## ACKNOWLEDGMENTS

The authors thank the CRCHUM BSL3 and Flow Cytometry Platforms for technical assistance, and Mario Legault for cohort coordination and clinical samples. Figures 1A, 6A and 8 were created using BioRender.com. We thank the following collaborators for kindly providing plasmids to produce antibodies: James Robinson (Tulane University) for A32 and 17b, the NIH AIDS Reagent Program for 2G12 and 246D.

Conceptualization: E.B., A.T. and A.F.; Methodology: E.B., A.T., G.T., M.C., D.Y. HC.C., J.R, WD.T., J.R., S.S., M.P. and A.F.; Investigation: E.B., A.T., F.T.,; Resources: M.D., D.M.H. M.P. and A.F.; Formal Analysis: E.B., A.T., F.T., S.S., and A.F. Supervision: M.D., D.M.H., S.S., M.P. and A.F.; Funding acquisition: M.P., A.F.; Writing – original draft: E.B. and A.F. Writing – review & editing: E.B., A.T., F.T., M.C., D.Y., HC.C., TJ.C., C.B., H.M., W.D.T., M.D., J.R., D.M.H., S.S., M.P. and A.F..

## DATA AVAILABILITY STATEMENT

Data and reagents are available upon request.

## ETHICS APPROVAL

Written informed consent was obtained from all study participants and research adhered to the ethical guidelines of CRCHUM and was reviewed and approved by the CRCHUM institutional review board ((Ethics Committee approval nos. 22.173, 11.063, MP-02-2024-11734 and 11.062)). Research adhered to the standards indicated by the Declaration of Helsinki. All participants were adult and provided informed written consent prior to enrolment in accordance with Institutional Review Board approval.

## DISCLAIMER

The views expressed in this manuscript are those of the authors and do not reflect the official policy or position of the Uniformed Services University, the US Army, the Department of War, the Henry M. Jackson Foundation for the Advancement of Military Medicine, Inc. or the US government. The funders had no role in study design, data collection and analysis, decision to publish, or preparation of the manuscript.

---

**Figure S1.**
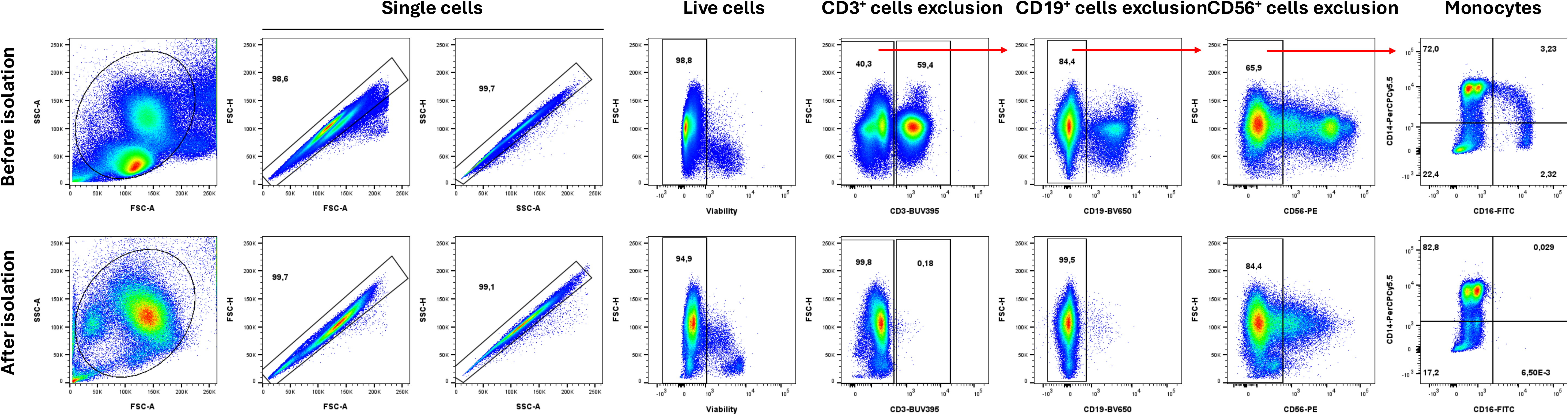
Flow cytometry analysis of purity and efficiency of monocyte isolation. Total PBMCs and isolated monocytes were stained to evaluated monocyte isolation efficiency. Cells were identified according to cell morphology by light-scatter parameters and exclusion of doublets. Monocytes were then identified after exclusion of CD3^+^, CD19^+^ and CD56^+^ cells.

**Figure S2:**
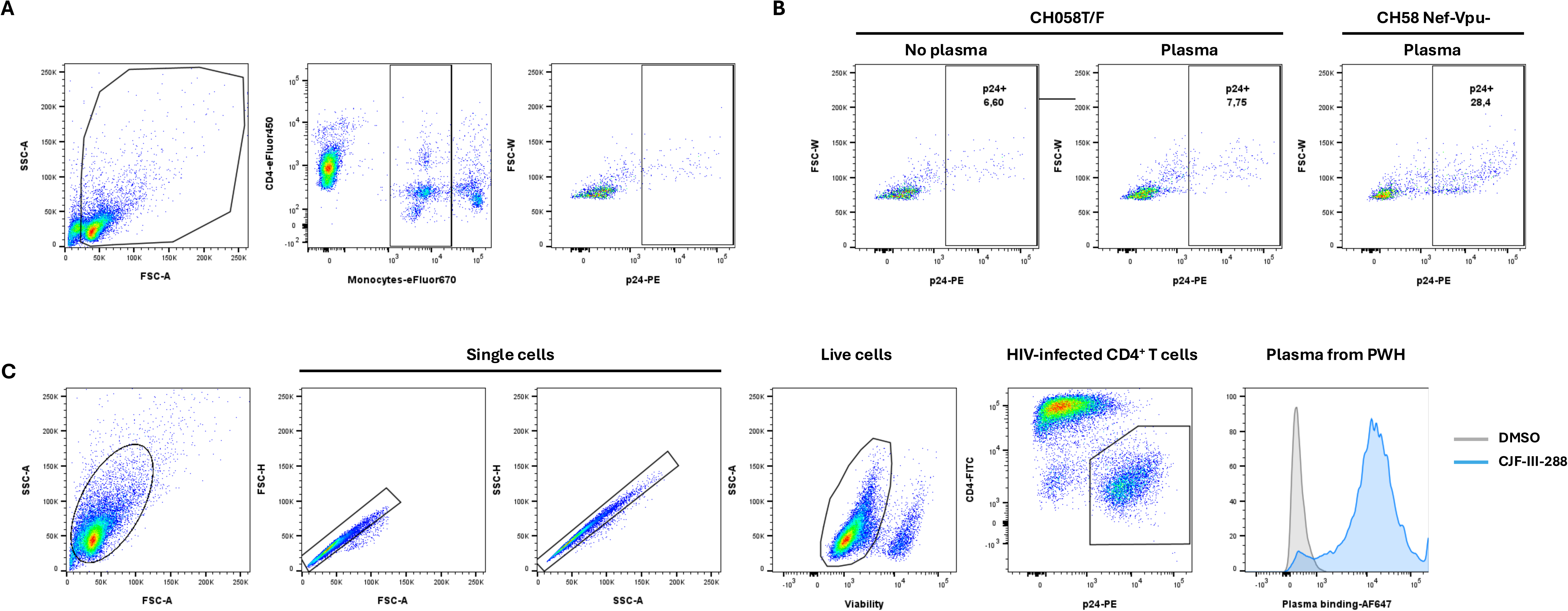
Gating strategy for ADCP assay and HIV-1 Env recognition by antibodies plasma from PLWH. (A) Representative flow cytometry gates to measure ADCP activity. Monocytes were identified according to cell morphology by light-scatter parameters (left) and CD4^+^ T cells (eFluor450^+^eFluor670^-^) exclusion (middle). Subsequent gating was done on the p24^+^ population on monocytes (right). (B) Representative plots for the p24^+^ gate for target CD4 T cells infected with HIV-1_CH058T/F_ WT (left and middle) or Nef-Vpu- (right) in absence (left) or in presence (middle and right) of plasma from PWH. (C) Representative flow cytometry gates to identify the level of Env recognition at the surface of HIV-1-infected CD4^+^ T cells by antibodies and plasma. Cells were identified according to cell morphology by light-scatter parameters and exclusion of doublets. Cells were then gated on living cells. Finally, the median of fluorescence of Alexa Fluor 647 (plasma binding level) was measured in infected p24^+^CD4^low^ cells.

**Figure S3.**
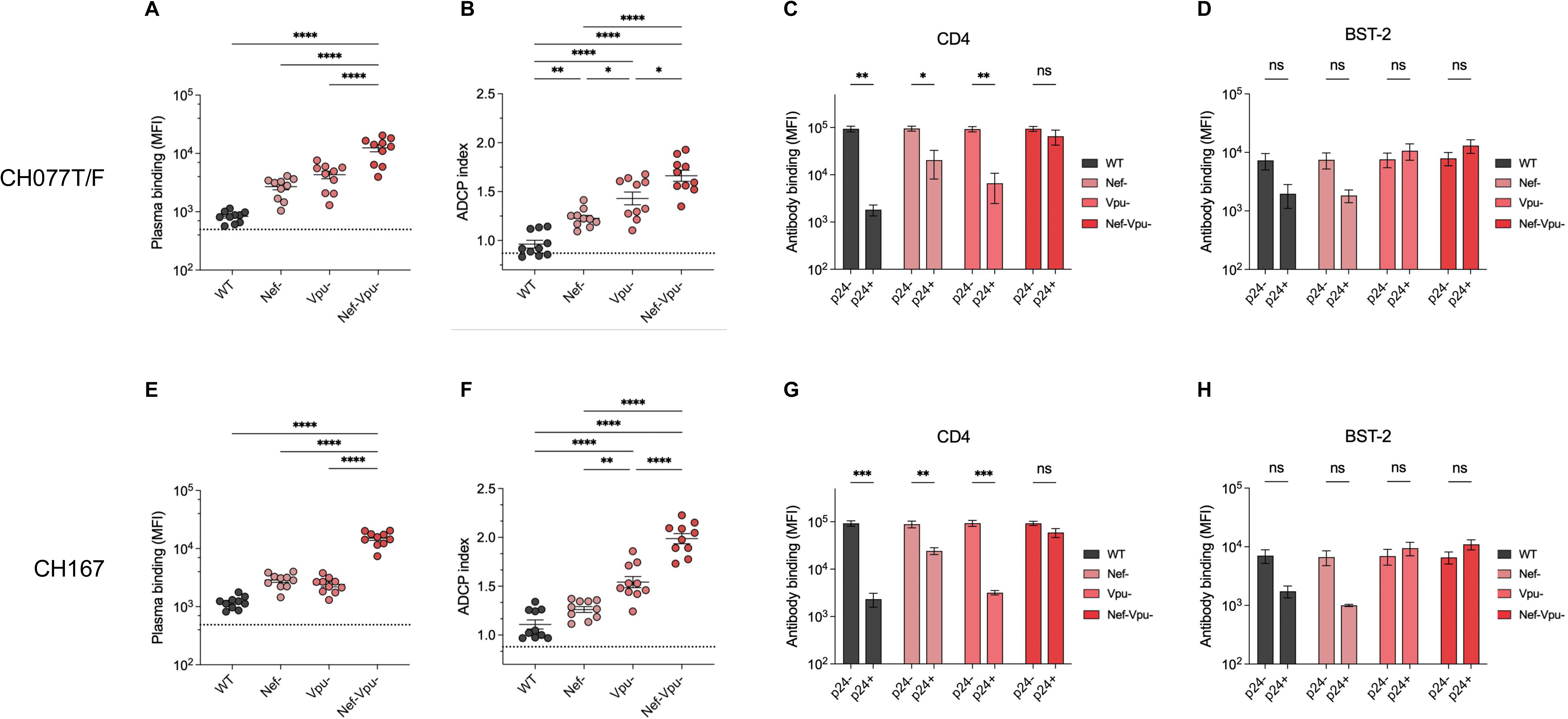
Effect of HIV-1 accessory proteins on recognition and ADCP-mediated killing of infected cells by PWH plasma. (**A**) Recognition and (**B**) PWH plasma-mediated ADCP killing of CD4^+^ T cells infected with HIV-1_CH077T/F_ harboring no mutation (WT), single deletion of the Nef (Nef-) or Vpu proteins (Vpu-) or deletion of both accessory proteins (Nef-Vpu-). Dotted lines represent the mean plasma binding or ADCP activity that was measured with 5 plasmas from PWoH. (**C**) Expression of CD4 and (**D**) BST-2 at the surface of p24^-^ or p24^+^ cells infected with the indicated HIV-1 isolates. (**E**) Recognition and (**F**) PWH plasma-mediated ADCP killing (**F**) of CD4^+^ T cells infected with HIV-1_CH167_ harboring no mutation (WT), single deletion of the Nef (Nef-) or Vpu proteins (Vpu-) or deletion of both accessory proteins (Nef-Vpu-). Dotted lines represent the mean plasma binding or ADCP activity that was measured with 5 plasmas from PWoH. Expression of (**G**) CD4 and (**H**) BST-2 at the surface of p24^-^ or p24^+^ cells infected with the indicated HIV-1 isolates. Statistical significance was determined by one-way ANOVA for **A-B-E-F** and two-way ANOVA for **C-D-G-H.**

**Figure S4.**
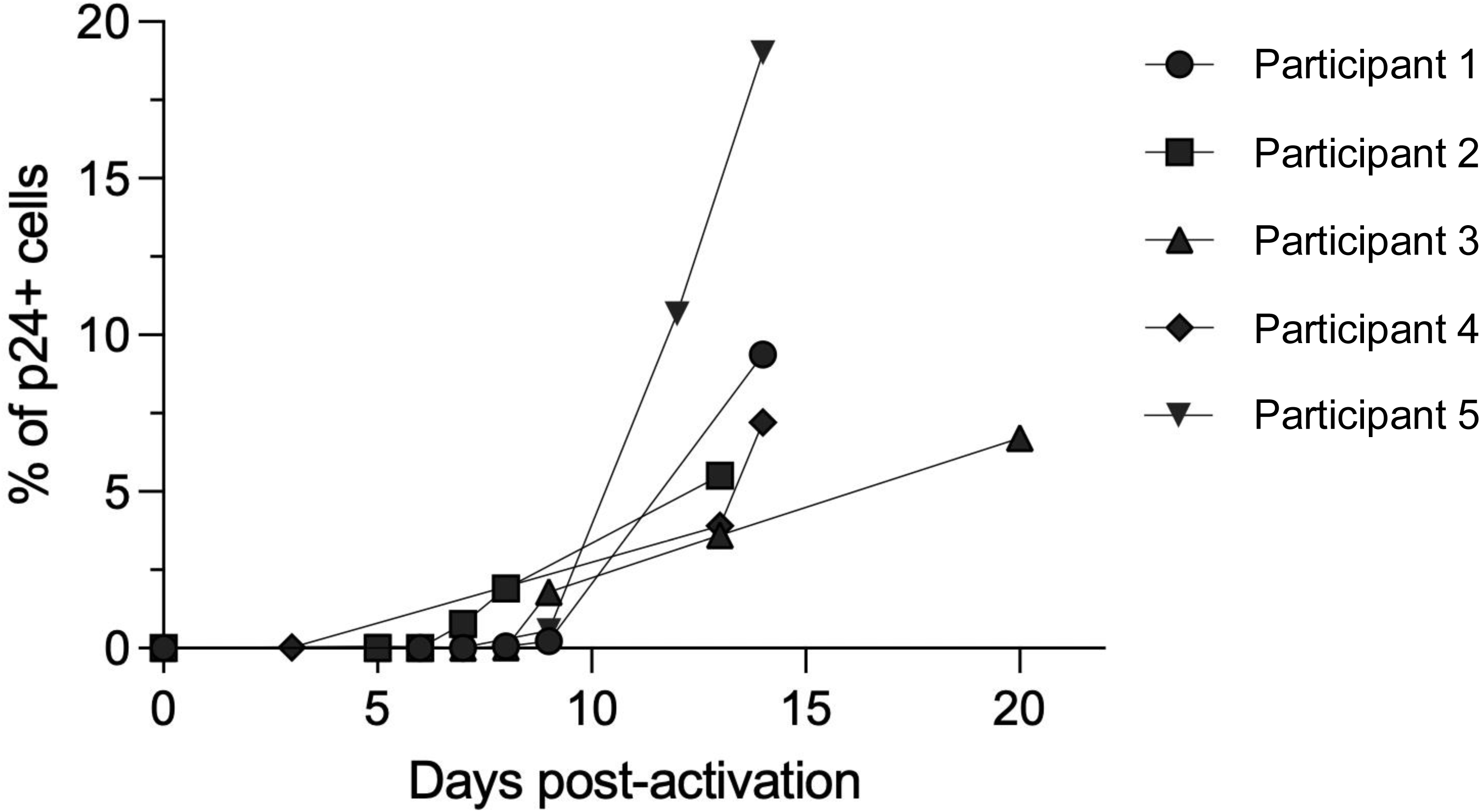
Expansion of *ex vivo* p24^+^ CD4^+^ T cells from PWH after reactivation. CD4^+^ T cells were isolated from the PBMCs of PWH under ART and were reactivated with phytohemagglutinin-L (10 µg/ml) for 48 hours. Cells were maintained in complete RPMI 1640 medium supplemented with 100 U/ml of IL-2 and p24 levels were monitored until they reached at least 5% of the CD4^+^ T cell population.

**Figure S5:**
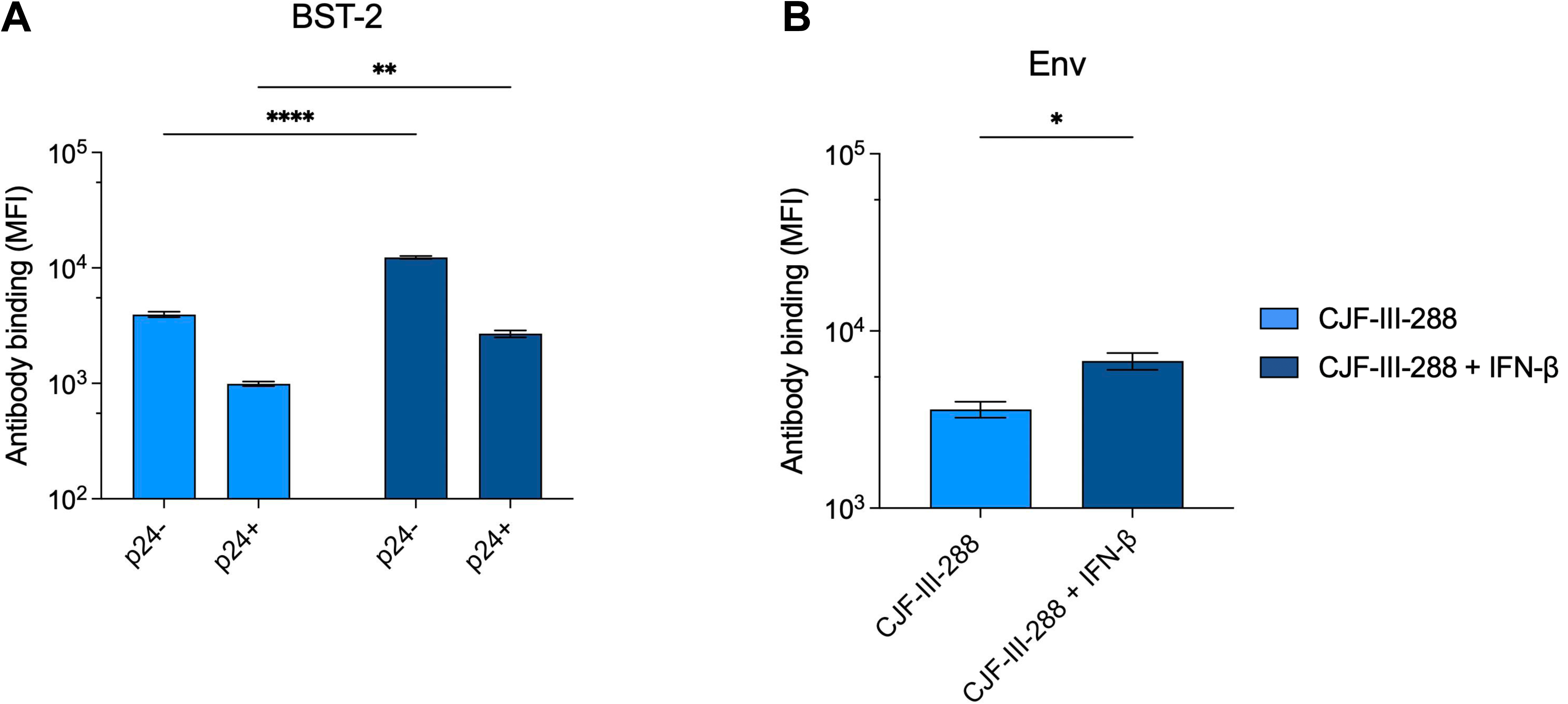
**Cell surface expression of BST-2 and Env after IFN-β treatment**. Primary CD4^+^ T cells were infected with HIV-1_CH058T/F_ for 48 hours and treated with IFN-β (2ng/mL) for 24h. Expression of (**A**) BST-2 at the surface of p24+ and p24-cells was measured with an anti-BST-2 antibody and (**B**) Env expression at the surface of p24+ cells was measured with the 2G12 antibody, and data was acquired by flow cytometry.

**Table S1.** Information on PWoH PBMC donors

| **Sex** |  | **Age** |
| --- | --- | --- |
| Female (n) | Male (n) |  |
| 2 | 6 | 49 (40 – 68) |

\*Values presented are medians with ranges in parentheses

**Table S2.** Information on PWH PBMC donors used for *ex vivo* CD4^+^ T cell expansion

| **Participant** | **Sex** | **Age** | **Time since acquisition (years)** | **Time since ART initiation (years)** | **Viral load (copies/mL)** |
| --- | --- | --- | --- | --- | --- |
| 1 | Male | 44 | 14 | 3 | Undetectable |
| 2 | Male | 59 | 34 | 2 | Undetectable |
| 3 | Male | 53 | 28 | 13 | Undetectable |
| 4 | Male | 61 | Unknown | 13 | Undetectable |
| 5 | Male | 54 | Unknown | 13 | Undetectable |

**Table S3.** Information on PWH plasma donors

| **Sex** |  | **Age** | **Years since**<br>**ART initiation** | **Viral load**<br>**(copies/mL)** | **CD4 count**<br>**(cells/mm<sup>3</sup>)** |
| --- | --- | --- | --- | --- | --- |
| Female (n) | Male (n) |  |  |  |  |
| 3 | 17 | 52 (40 – 69) | 18.2 (0.4 – 23) | Undetectable | 677 (197 – 1250) |

\*Values presented are medians with ranges in parentheses

